# Sysrev: A FAIR platform for Data Curation and Systematic Evidence Review

**DOI:** 10.1101/2021.03.24.436697

**Authors:** Thomas Bozada, James Borden, Jeffrey Workman, Mardo Del Cid, Jennifer Malinowski, Thomas Luechtefeld

## Abstract

**Abstract:** Well-curated datasets are essential to evidence based decision making and to the integration of artificial intelligence with human reasoning across disciplines. However, many sources of data remain siloed, unstructured, and/or unavailable for complementary and secondary research. Sysrev was developed to address these issues. First, Sysrev was built to aid in systematic evidence reviews (SER), where digital documents are evaluated according to a well defined process, and where Sysrev provides an easy to access, publicly available and free platform for collaborating in SER projects. Secondly, Sysrev addresses the issue of unstructured, siloed, and inaccessible data in the context of generalized data extraction, where human and machine learning algorithms are combined to extract insights and evidence for better decision making across disciplines. Sysrev uses FAIR - Findability, Accessibility, Interoperability, and Reuse of digital assets - as primary principles in design. Sysrev was developed primarily because of an observed need to reduce redundancy, reduce inefficient use of human time and increase the impact of evidence based decision making. This publication is an introduction to Sysrev as a novel technology, with an overview of the features, motivations and use cases of the tool.

**Methods:** Sysrev.com is a FAIR motivated web platform for data curation and SER. Sysrev allows users to create data curation projects called “sysrevs” wherein users upload documents, define review tasks, recruit reviewers, perform review tasks, and automate review tasks.

**Conclusion:** Sysrev is a web application designed to facilitate data curation and SERs. Thousands of publicly accessible Sysrev projects have been created, accommodating research in a wide variety of disciplines. Described use cases include data curation, managed reviews, and SERs.

---

A systematic evidence review (SER) is a methodological tool used in a variety of disciplines to capture all relevant published literature on a topic, which may lead to quantitative analysis of the data (meta-analysis) or qualitative synthesis. Core principles of SERs are rigor, reproducibility, and transparency. SERs generally follow a prescribed process, beginning with the identification of the population, intervention, comparator, and outcomes of interest. Once the relevant key questions have been developed, a search in one or more databases yields a number of articles that must be reviewed, typically in duplicate by blinded reviewers according to pre-specified inclusion and exclusion criteria. Data from included studies are abstracted into forms for subsequent quantitative analysis or qualitative synthesis.

One of the primary goals of Sysrev is to apply SER methods to other workflows, including generalized data curation, education, and any workflow involving collaboration between humans and between humans and machines for evaluation of digital documents.

Although SERs accompanied by meta-analysis are considered the highest level of evidence, due to their overall objectivity, they are time-consuming [1]. Adhering to SER methodology can be difficult, particularly when the method is applied to non-biomedical research [2]. Thorough searches can require subscriptions to multiple databases. Depending on the topic, it is not unusual for a well-designed literature search to return more than 10,000 studies and for a SER to take more than one year to complete. To alleviate some of the inherent challenges with managing a large corpus of literature, the number of participants on a review may be quite large. Numerous software have been developed to help reviewers manage the literature base and perform one or more phases of a SER, such as citation management or title and abstract screening [3]. More recently, machine learning algorithms have been proposed as a way to increase workload efficiency [4]. However, despite the plethora of available SER software, recent reviews have found few software can adeptly meet the needs of those who perform SERs across disciplines [5].

While the number of SER publications has risen substantially in the past decade, much of the data extracted from their included articles remains locked away, limiting their reuse by others performing adjacent research. For example, data extraction templates may be published as a supplement to a SER, but the completed extraction forms for every study may not, or what is published is in a form that precludes secondary research. Therefore, data curation is an essential component of data re-use and research transparency/reproducibility.

There are a number of curated, publicly-available (open access) data repositories for scientific and academic disciplines, such as the worldwide Protein Data Bank (wwpdb.org), the Cambridge Structural Database (https://www.ccdc.cam.ac.uk/solutions/csd-core/components/csd/), and the Publishing Network for Geoscientific & Environmental Data (PANGAEA) (https://www.pangaea.de/). However, re-use of the information can be limited due to controls on database access, limited labeling of semi-structured or unstructured data [6], and the structures of databases that prevent inclusion of some experimental data [7], among others. These obstacles contribute to “dark data” the data from small experiments which are unlikely to be openly shared with other researchers and exist only on controlled-access laboratory or university servers [8]. Consequently, much of publicly funded research remains relatively hidden and at risk for loss unless submitted to repositories, disproportionately affecting the ‘long tail’ of science and technology [9].

The FAIR Guiding Principles were published to encourage better data stewardship and management and in support of infrastructure to minimize dark data [7]. The FAIR Principles refer to Findability, Accessibility, Interoperability, and Reuse of digital data. The findability tenet incorporates the assignment of unique identifiers and metadata to information so that it can be found more easily by humans or machines (especially search engines). Accessibility requires the data or metadata be retrievable using a standard communications protocol and that the metadata associated with the data remain available even after the original data is no longer accessible. Accessibility does not require all types of data be freely available to any user for any purpose; for example, the National Center for Biotechnology Information’s dbGaP repository of phenotype and genomic information restricts access to datasets to NIH and senior investigators [10]. Incorporation of shared and standardized vocabularies are essential to support interoperability and reuse of data and its metadata. Importantly, FAIR Principles are discipline agnostic; that is, they can be applied across a range of academic and industrial activities. Since their 2016 publication, numerous disciplines are striving to apply FAIR Principles, including genomics, agriculture [11, 12], climate science, and medicine.

Sysrev, which is in active development, is a FAIR motivated software to extract and label data from a variety of data sources. This paper describes the features behind Sysrev, provides use case examples, gives a brief meta-analysis of Sysrev open access data, and suggests future directions and enhancements to Sysrev.

## Workflow

The following section describes the user experience and project workflow within Sysrev.

First, Users access and create accounts at sysrev.com. All account types (including free accounts) can create an unlimited number of publicly accessible Sysrev projects. Public projects can be discovered by search engines like google and bing. All data generated in a public project can be discovered and downloaded on the internet.

Users with paid accounts can choose to designate any of their Sysrev projects to remain private. Private project data can only be accessed by project members.

Sysrev projects can be broken down into 6 phases: data source creation, label definition, user recruitment, review, data export, and analysis (Figure 1).

**Figure 1.**
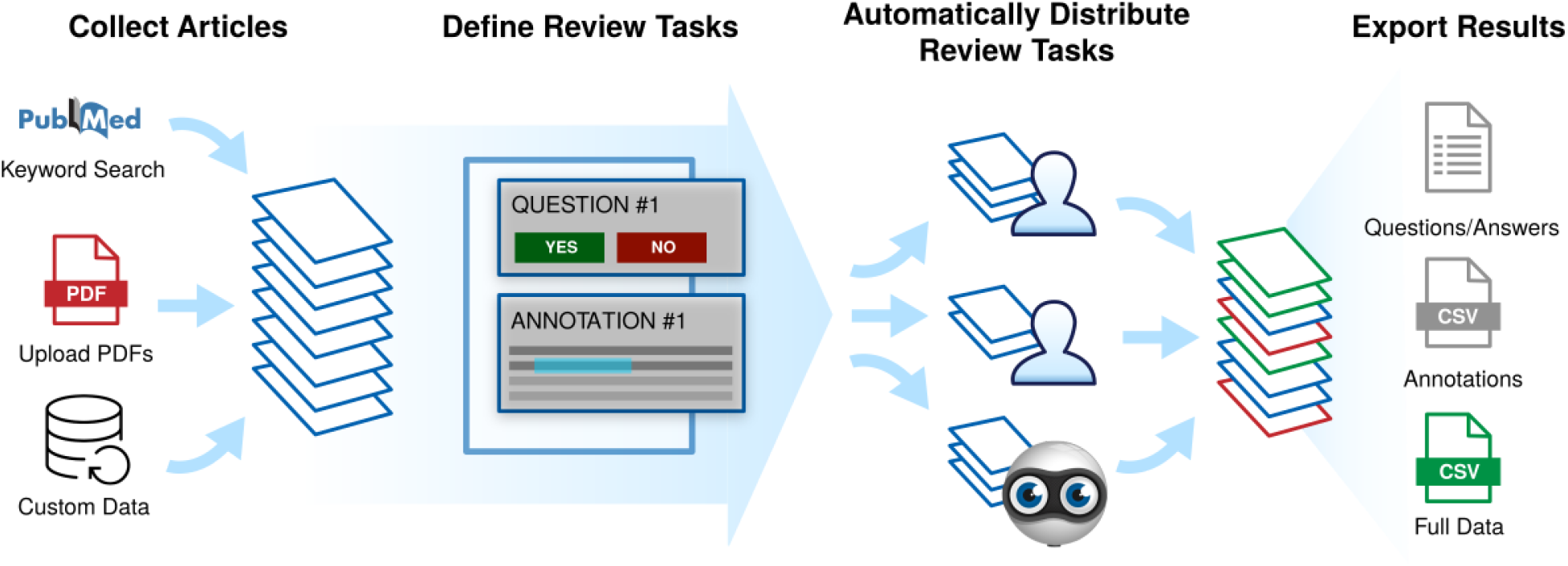
Sysrev projects can be broken into 6 stages from left to right (1) articles are collected to form a data source (2) review tasks or ‘labels’ are defined (3,4) reviewers are recruited and asked to complete review tasks. Active learning is involved at this stage to prioritize and replicate labelling tasks (5,6) data is exported and analyzed.

### Data Source Creation

Sysrev projects are highly customizable and can accommodate a variety of data import structures. Currently supported file imports include XML, RIS citation format, and PDF. Text, JSON, and HTML formats are also supported via the Sysrev programming interface. A growing number of data integrations including pubmed.gov and clinicaltrials.gov searches.

#### Future development

Sysrev is built to generalize the systematic review process to review any kind of digital document. Future sources will include support for images, videos, and other custom formats that can be viewed and annotated by users in a browser.

Future Sysrev sources will allow the use of one sysrev as a source for another sysrev, thus facilitating stagewise reviews and other applications involving review of reviews. Living reviews will be enabled by allowing sources to grow over time. Source publication will enable users to develop their own sources which can be shared between sysrevs. Sysrev already supports programmatic source publication at https://datasource.insilica.co, but these features are not yet integrated into the Sysrev user interface.

### Label Definition

Labels are structured data elements that are extracted from documents. Sysrev supports basic labels, group labels, and is developing many more advanced label types.

Sysrev supports three basic label types: boolean, categorical and string labels. Label options allow labels to be set as “required”, “requiring user consensus”, as part of inclusion criteria, or set with a variety of other options.

“Required” labels must be provided by reviewers, reviewers cannot submit a document review until all required labels have been provided.

Labels that require user consensus will mark documents as “in conflict” when different reviewers disagree on the value associated with the label in a given document, this is a method for identifying documents with inconsistent user labels.

Labels with inclusion criteria rules are associated with document screening. Inclusion criteria options allow administrators to capture the reasons that a document was included or excluded.

Among a variety of other options, String labels can have associated regular expressions. Regular expressions allow users to write a pattern that must be met by user inputs. Regular expressions can be used to modify string labels to behave as numeric labels, date labels, dosage labels, or anything else that can be encoded in a regular expression, which is almost any pattern that can be written in text. Sysrev strongly recommends using regular expressions to normalize the string inputs that users can provide.

Group labels allow project administrators to extract table data from documents. Group labels are composed of a set of basic labels. When extracting data into a group label, reviewers fill in rows in a table corresponding to the provided basic labels. Figure 2 provides a screenshot where a reviewer is extracting testing data from a safety data sheet [13]. In this example, the project administrator wanted to know what test types were defined in the safety data sheet, and to associate related dose amounts, results, species, etc. Group labels are a powerful concept for creating relational databases from documents. When they are exported, each group label can be treated as a new table.

**Figure 2.**
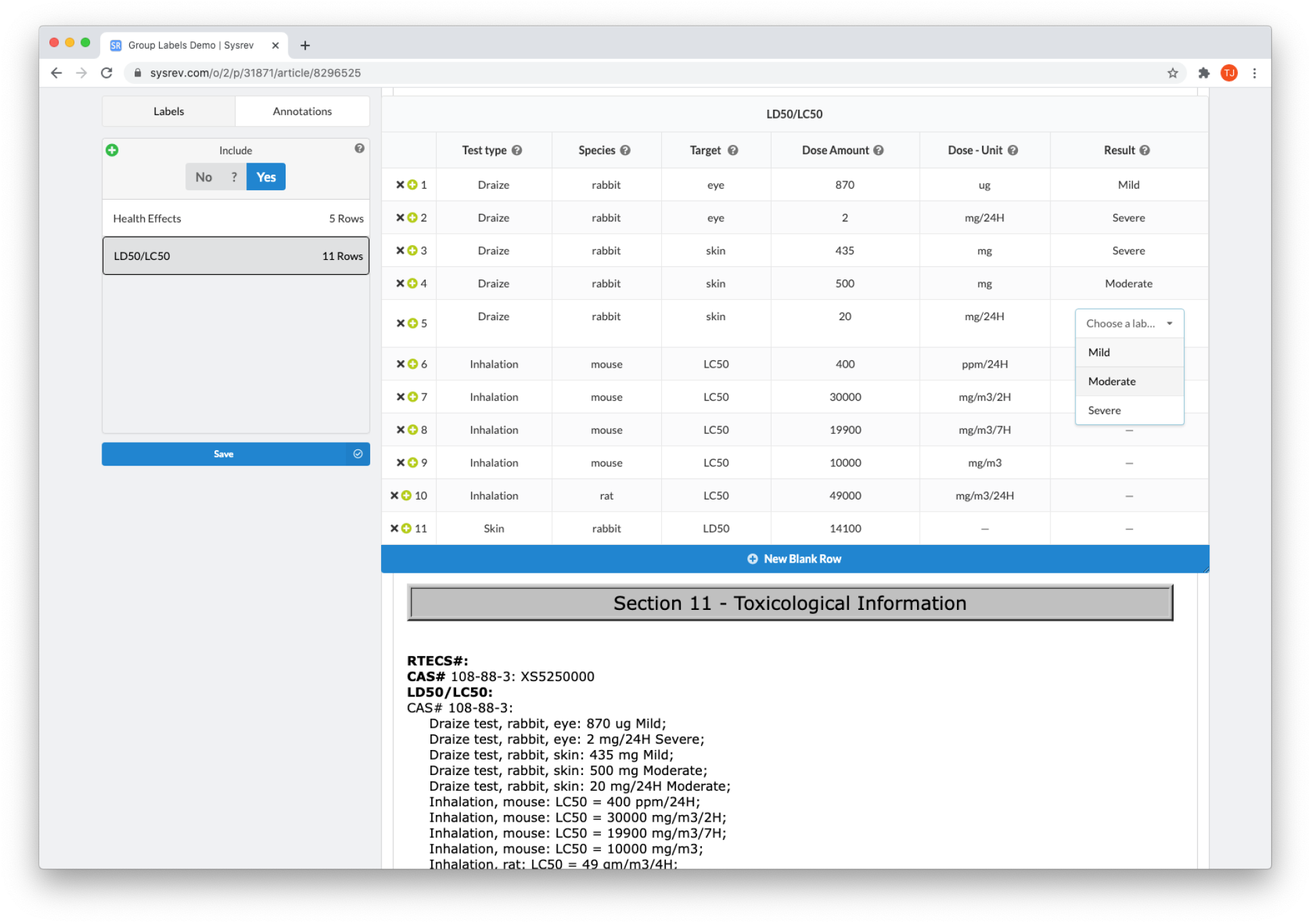
Users extract testing information from a safety data sheet using a group label. This example is available at sysrev.com/o/2/p/31871/article/8296525.

Label definitions are a generalizable concept. When they are built correctly, they communicate to reviewers specific constraints for extracting data from documents. They can even involve interactions with documents. For example, Sysrev supports “text annotation” which involves the selection of text within a document and the assignment of a value to the selection. Sysrev supports this advanced kind of text annotation label.

#### Future Development

Other advanced label types and label properties are in development, such as image annotation, shared labels which can be shared across many projects, and question/answer labels which will allow sysrev to function as an educational platform.

### User Recruitment

After creating document sources and defining labels, users must be recruited to begin the reviewing process. Many sysrev projects consist of a single project owner who also performs the full review. Other projects have hundreds of reviewers. Most fall somewhere in-between. Recruiting users can be done by sharing a link or sending emails directly from sysrev.com.

Sysrev also supports recruitment of existing Sysrev users using sysrev.com/users. Users can identify themselves as available for future reviews. Before inviting these users, project administrators can review previous user projects and performance by using sysrev.com/search or viewing the user profile page.

Users can be assigned to groups. Eventually these groups will allow project administrators to modify review prioritization and tasks for specific users. For example, reviewers could be grouped by language and assigned articles accordingly. Or, biology experts could be asked to perform one task, while chemistry experts are asked to perform a different task on the same documents.

User recruitment is one area where Sysrev is attempting to improve the visibility and community behind review work. Some Sysrev users have performed tens of thousands of document reviews, their experience and publicly visible work could make them strong candidates to help in other projects.

Document review is the heart of Sysrev. Sysrev distributes documents to reviewers and manages that distribution according to several project parameters. Projects can require double review, single review, or unlimited review (where all reviewers review all documents). Review can be prioritized in order to maximize the rate of document review, in order to maximize the rate of review duplication, or to balance these priorities.

Unlimited review can be useful for classroom applications where students are evaluated according to some gold standard values associated with each document.

#### Progression

Project administrators can check in on their project at any time to keep track of completion and reviewer performance. Sysrev analytics features allow administrators to evaluate reviewer performance relative to other users. The distribution of extracted labels, and other informative analytics are provided on the project overview and analytics pages, and are updated in real time during review. Figure 3 shows the Sysrev label counting tool, which provides a visualization and filtering options to keep track of label distributions during project progression.

**Figure 3.**
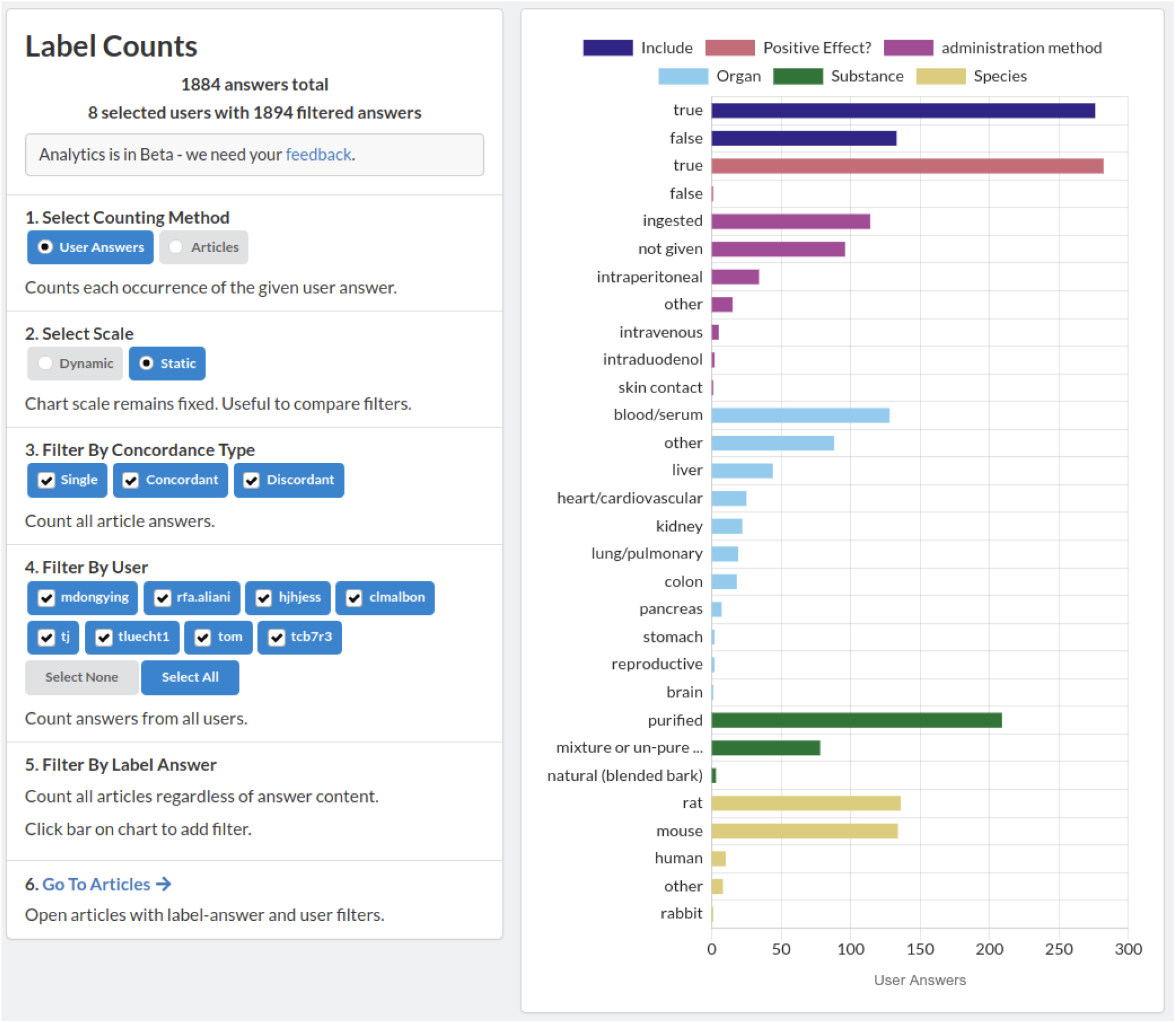
A screenshot of the sysrev label counting tool. A demonstration of this tool is publicly available at https://sysrev.com/o/2/p/21696/analytics/labels. Label counts can be filtered, visualized and updated in real time as a sysrev progresses.

#### Active learning

During the review process, Sysrev periodically creates machine learning models. These models attempt to replicate all binary and categorical review tasks. Models are also used to prioritize review, by selecting documents with the lowest classification confidence in an active learning paradigm [14]. Anecdotally, when active learning was implemented on Sysrev, users reported that the more difficult documents in a review were quickly being prioritized. The motivation behind Sysrev’s active learning is to maximize the value of human time. When machine learning models can accurately evaluate uncertainty, they can be used to prioritize human review to only those documents that are difficult to automate.

#### Conflict Resolution

A conflict flag is generated when users disagree about labels marked as “requiring concordance” in a specific document. Conflict flags identify documents where reviewers disagreed about one or more labels that require concordance. Once an article is flagged for conflict, the conflict can be resolved by a project administrator. By default, the inclusion label, which is used to screen articles in a sysrev, requires concordance.

### Data Export

Sysrev supports methods to export review data to spreadsheets, xml documents, and json files. In addition, a graphql API is available that allows users to programmatically access sysrev data. Two open source clients of this graphql API written in R and Python are in development at github.com/sysrev/RSysrev and github.com/sysrev/PySysrev.

### Analysis

Sysrev provides a growing set of analytics tools built directly into each project.

#### Concordance

The Sysrev concordance tool allows project administrators to evaluate their labels and reviewers. Concordance is measured by counting how often users agree about extracted labels. Figure 4 demonstrates the concordance tool in a public review on mangiferin (https://sysrev.com/p/21696/analytics/concordance).

**Figure 4.**
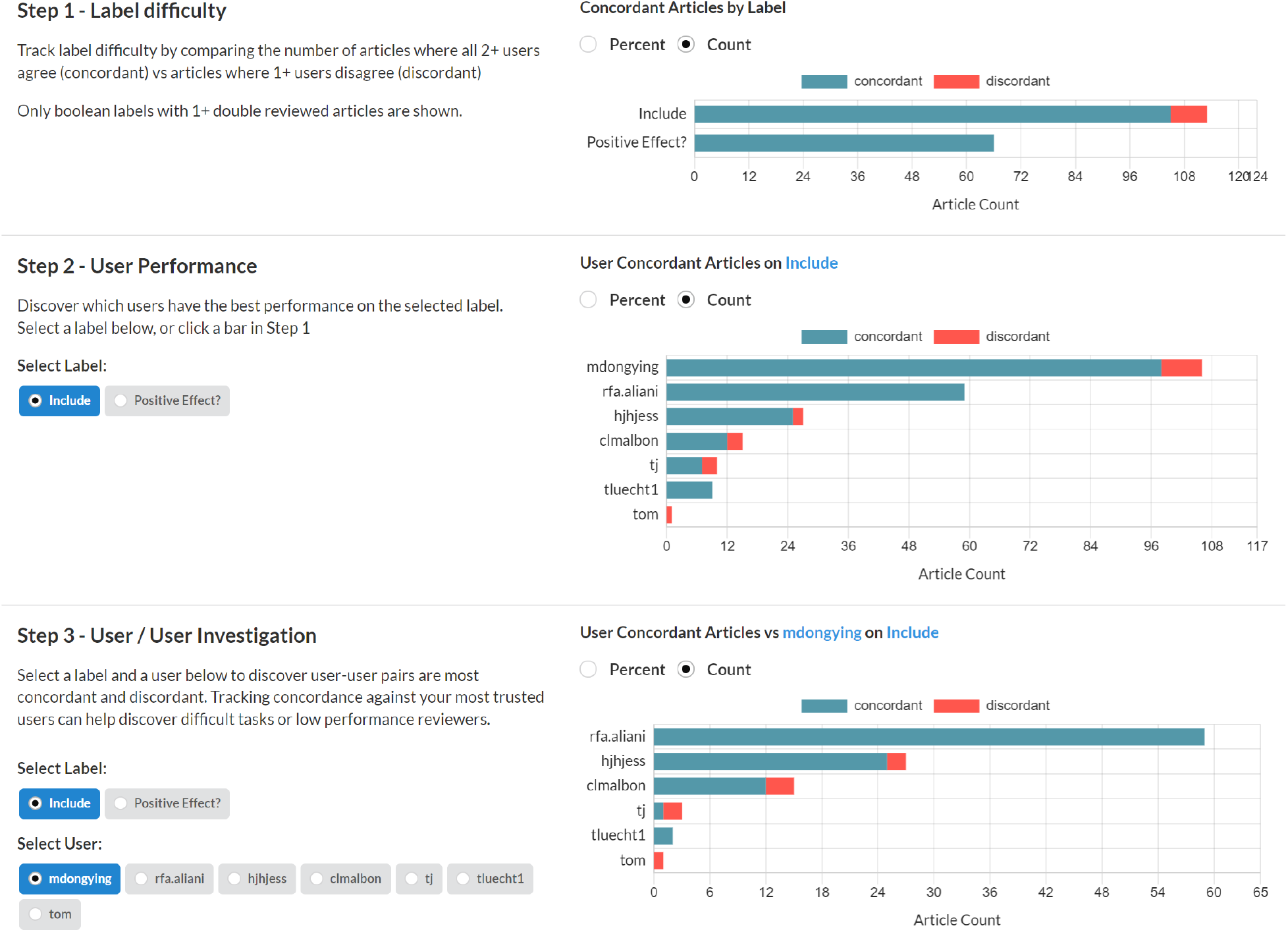
A screenshot of the Sysrev concordance tool in action. This tool can be demonstrated at sysrev.com/p/21696/analytics/concordance

In the first step, administrators can quickly evaluate which tasks are most concordant. Highly concordant tasks are tasks that reviewers frequently agree on, and are more likely to be well defined and understood tasks. Highly discordant tasks may indicate areas where reviewers have been inadequately trained or tasks have been underdeveloped.

In the second step administrators can view how concordant each user is on a given label. In Figure 4, User rfa.aliani has perfect concordance on the include label, and other users have varying levels of discordance.

In the last step, administrators view users’ concordance against a given user for a certain label. Figure 4 demonstrates that rfa.aliani has perfect concordance with mdongying for the include label, but that other users have some discordance with mdongying.

### Machine Learning

Sysrev periodically builds machine learning algorithms that attempt to replicate all boolean, categorical and (soon) annotation reviewer tasks. Sysrev provides some transparency around the accuracy for these models via a histogram describing model predictions for articles that human reviewers have already excluded and included. Machine learning models are updated periodically as articles are screened and new histograms are provided to evaluate model performance [15]. Figure 5A provides several example histograms, from public Sysrev projects, which are automatically generated during project progression. These histograms show the number of articles included or excluded by human reviewers with a probability of inclusion given on the x axis.

**Figure 5.**
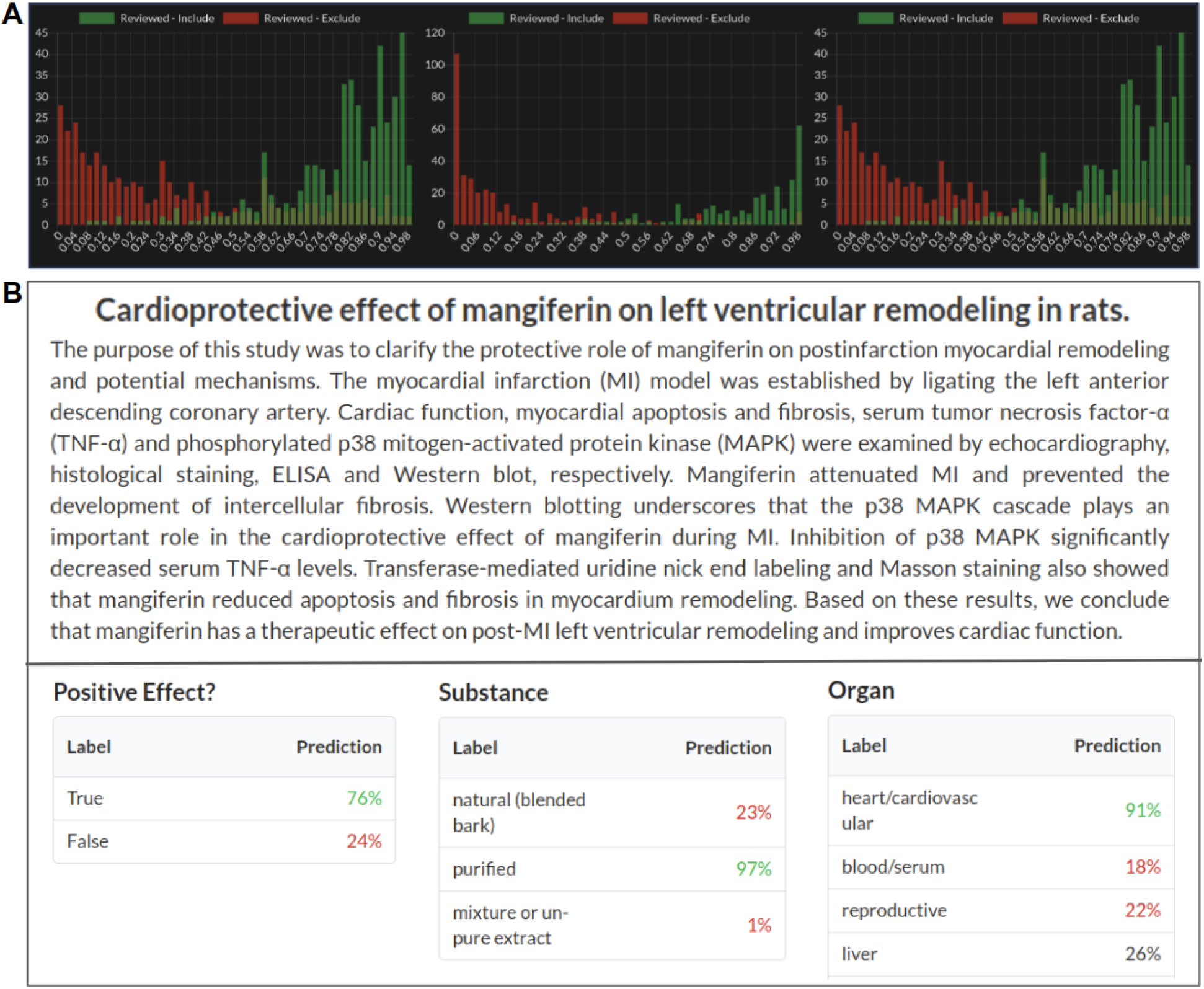
**A**. Machine learning prediction of article inclusion compared to actual reviewer inclusion decisions from 3 public sysrev projects on spinal surgery sysrev.com/p/14872, sysrev.com/p/14873, sysrev.com/p/14874. **B.** Sysrev categorical and boolean predictions. Sysrev predictions correctly identify that this article references a “positive effect” on “heart/cardiovascular”. They provide a strong prediction of “purified” mangiferin substance, even though the article does not explicitly specify this. Model predictions can be seen on every article in a project. This screenshot comes from an Insilica.co project on Mangiferin (sysrev.com/p/24557/article/7225450).

Sysrev automatically builds models to replicate every categorical and boolean label in every sysrev project. As of March 1, 2021 a total of 94,064,632 predictions have been made by Sysrev models for articles in sysrevs. The predictions for these labels can be viewed in articles and used to filter article lists. For example, after labeling a few hundred articles for a boolean “heart/cardiovascular” label, the resulting model can be used to filter articles with a high probability of receiving the “heart/cardiovascular” label.

Figure 5B shows label predictions for 3 of the categorical / boolean labels for an article in a sysrev on Mangiferin, a mango extract. This review has many categorical and boolean variables, three of which are:

**Positive effect** - Is the discussed effect a positive or negative one?
**Substance** - What form of mangiferin was used? Natural, purified or mixture
**Organ** - What organ(s) were referenced: Heart, blood, reproductive, liver, etc.

Sysrev models provide a value between 0 and 1 (interpreted as a percentage) for each label + answer pair. Boolean labels receive two predictions, one for label + true and one for label + false. Categorical labels receive a prediction for each label + answer, these values do not necessarily sum to 1, although future features will update model predictions to normalized values for single answer (mutually exclusive) categoricals.

## Supplemental Features

Sysrev develops a growing set of features that are supplementary to the core workflow features.

### Payment Platform

Sysrev provides a payment platform where project administrators can deposit and distribute funds to reviewers on a per-document reviewed basis. In the future, Sysrev’s payment platform will integrate modeling, previous user performance, and the quality/difficulty of document reviews into rewards. We hope to provide a marketplace to link data curation projects to reviewers.

### Project Cloning

One unique aspect of Sysrev is the ability to clone projects. This provides researchers working on related sysrevs, others who want to use the same data templates, or those who want to work with shared data, an opportunity to streamline project setup and ensure consistency in how data is extracted/curated across projects. Cloned sysrevs maintains the same labels and structure as the initial review, ensuring reproducibility. One example of a cloned project is a SER of methods used to quantify the mechanical work of breathing during exercise (available at: sysrev.com/p/27698). After the initial review was completed, the authors cloned the existing project to update the literature search (see sysrev.com/p/50226). Project cloning is motivated by analogy between sysrev projects and git repositories, we aim to make sysrev projects easier to copy and facilitate the dissemination of review protocols.

These are the major features behind a Sysrev workflow. The Sysrev development roadmap is driven by user requests. Please contact us at with any requests.

## Case Studies

We invited early Sysrev users to describe their own projects. These case studies serve as a demonstration of several possible use cases for sysrev.

### Ascorbate Administration Methods in Oncology Clinical Trials[16]

**Author:** Channing J Paller^1^, Tami Tamashiro^2^, Thomas Luechtefeld^3^, Amy Gravell^2^, Mark Levine^4^

1. Johns Hopkins School of Medicine
2. EMMES
3. Insilica LLC
4. National Institute of Health/National Institute of Diabetes and Digestive and Kidney Diseases

**Sysrev:** https://sysrev.com/p/6737 - Vitamin C Cancer Trials

#### Description

Vitamin C (ascorbic acid, ascorbate) is an essential micronutrient and versatile reducing agent found in food and dietary supplements. While vitamin C can reduce reactive oxygen species at micromolar concentrations functioning as an antioxidant, it can also act as a pro-oxidant at higher millimolar concentrations, killing cancer cells in vitro and slowing in vivo tumor growth.

A review was performed of all clinical trial data from trials submitted to clinicaltrials.gov matching condition “cancer” and intervention “ascorbic acid”, “ascorbate”, or “vitamin c”. Information regarding administration methods (dosing, oral vs IV, multivitamin delivery), trial parameters (blinding, placebo, and randomization methods), drug combinations and disease types was extracted and organized for analysis. Analysis identified increased use of ascorbate alone and in combination with other therapies across multiple diseases. A high resolution evaluation of these outcomes is available as a book chapter in “Vitamin C and Cancer - An Overview of Recent Clinical Trials”[16]

### Data Curation Of Chemical Safety Data Sheets

**Author:** Daniel Mcgee, Wendy Schlett and Brad Van Valkenburg at Foresight Management

**Sysrev** https://sysrev.com/p/4047 many private reviews

#### Description

Safety Data Sheets (SDS) are documents containing the properties and occupational health, environmental hazards, and safety information for chemicals. SDS follow a consistent framework to align with the United Nations’ Globally Harmonized System of Classification and Labelling of Chemicals (GHS) and are required to be available to downstream users, who may include manufacturers or transporters. Chemicals of concern vary according to the purpose and for each company. For example, the European Chemicals Agency restricts the manufacturing, marketing, or use of substances under the Registration, Evaluation, Authorisation, and Restriction of Chemicals (REACH) Regulation while the California Office of Environmental Health Hazard Assessment maintains a list of chemicals known to cause cancer, birth defects, or other reproductive harm (Proposition 65) [17].

Sysrev was used by the Sustainable Research Group (SRG, now Foresight Management) to automate the extraction of relevant information from SDS to build a chemical of concern database for SRG clients. In addition to data from the SDS, SRG could label data with the name of the downstream user (account) or product, enabling cross-referencing to other chemical databases. This process facilitated attainment of LEVEL certification, an environmental certification for furniture similar to LEED certification for buildings, for a client [18].

### Methods Used To Quantify The Mechanical Work Of Breathing During Exercise

**Author:** Troy J Cross^1^, JW Duke^2^

1. Mayo Clinic & The University of Sydney, Australia
2. Northern Arizona University, USA

**Sysrevs**

https://sysrev.com/p/24827 & https://sysrev.com/p/46939 - Title & Abstract screening
https://sysrev.com/p/25285 & https://sysrev.com/p/49357 - Full-text screening
https://sysrev.com/p/27698 & https://sysrev.com/p/50226 - Data extraction

#### Description

The overarching purpose of this project is to survey the available literature for the many different methods used to quantify the mechanical work of breathing (Wb) during exercise. Each method of quantifying Wb has its own advantages and disadvantages, and their own technical requirements. Given the variety of methods used to measure Wb, we thought it would be prudent to systematically evaluate the literature and determine the relative popularity of each method, and to assess the quality of their implementation. By conducting this systematic review, we hope to raise awareness of the advantages/pitfalls associated with each method of quantifying Wb, and to emphasize the importance of reproducibility through clear descriptions of such methods. In doing so, we anticipate that these methods of quantifying Wb during exercise become more accessible to other Investigators in the area wishing to delve into the field of applied respiratory mechanics.

### SER: COVID-19 And Kidney Transplantation

**Author:** Ciara Keenan at Campbell UK & Ireland, Queen’s University, Belfast

**Sysrevs**

https://sysrev.com/p/29506 - COVID19 CKD
https://sysrev.com/p/30488 - COVID19 CKD ROBINS-I

#### Description

Numerous SERs have been performed using the Sysrev platform for one or more of the SER phases (screening, full-text review, and/or data extraction). An international team of researchers performed a SER using Sysrev to investigate the risks of COVID-19 to kidney transplant recipients [19, 20]. The reviewers assessed both English- and Chinese-language articles, uploading the studies as two separate .xml files. Sysrev allows users to sort results based on the import file; the team of reviewers used this function to review only the abstracts in their primary language.

A separate Sysrev project was created to manage the data extraction phase of the SER. As with article screening, the reviewers were able to sort the included studies by import file. The extraction form consisted of twenty-nine questions consisting mainly of string and categorical labels (e.g., age, health status). For the 31 included articles, more than 700 pieces of information were extracted, including the risk of bias for each study.

A third Sysrev project was created for the team to determine the risk of bias for each included study in the SER (available at: https://sysrev.com/u/249/p/30488). The Risk of Bias in Non-randomized Studies of Interventions (ROBINS-I) assesses risk of bias that may occur pre-intervention, at intervention, and post-intervention for a study. Domains evaluated include confounding, selection of participants into the study, classification of interventions, deviations from the intended interventions, missing data, measurement of outcomes, and selection of reported results. Each domain is individually examined, and an overall risk of bias is determined [21]. The authors created categorical labels for each of the ROBINS-I domains and an overall risk of bias.

### Wildlife Trade and Human Disease

**Author:** Hernan Caceres-Escobar^1^, Ciara Keenan^2^, Jon Paul Rodriguez^3^, Richard Kock^1^

1. Royal Veterinary College, University of London
2. Campbell UK & Ireland, Queen’s University, Belfast
3. IUCN Species Survival Commission

**Sysrev:** https://sysrev.com/p/43801 - Wildlife Trade

#### Description

The COVID-19 pandemic caused by SARS-CoV-2 is the latest in a series of novel diseases to affect humanity over recent decades. This has been counter to the trend of reduction in infectious disease burden enjoyed by humanity over the 20th Century due to the explosion in medical technologies and preventive measures. Initial circumstantial evidence suggested SARS-CoV-2 might have spilled into humans in a live wildlife trade market (commonly known as ‘wet markets’) in the city of Wuhan (Hubei Province, China), where multiple domestic and wildlife species and products were processed and sold.

This is not the first time human infectious diseases have been associated to the diverse trade in wildlife species, some proven such as the 2003 monkeypox outbreak (US), others unproven such as the 2002-2004 SARS outbreak (China) and COVID-19 index cases associated with Wuhan Market, China. Other human cases such as Ebola Virus infection were traced on a few occasions to index patients in contact with dead primates, great apes, and other wildlife species hunted and traded for food. The origins of HIV/AIDS are thought to be from evolution of simian (primate) origin virus (SIV) and the final example is during the bird flu pandemic (H5N1) in 2006, where a small number of imported birds into Europe showed infection with this virus, which led to a ban, having identified this wildlife trade hazard. Overall, zoonosis from wildlife trade appears to be rare (low risk or underreported), but as the examples illustrate, on occasions are of considerable public health concern. Given the seriousness of emerging pathogens, which have correctly or incorrectly been associated with wildlife trade, multiple organisations are now calling for bans on wildlife trade (i.e., from blanket bans to more nuanced and species-specific regulations). Wildlife trade has, despite the lack of hard evidence, been singled out as one of the main drivers of novel human disease emergence [22, 23]. In truth, the role of the wildlife trade in these events and zoonosis risks other than those mentioned above, remain poorly understood and largely undocumented.

In this project [24], we will identify and map all existing research evidence that primarily focuses on potential links between wildlife trade and human diseases (emerging human pathogens and zoonoses) and secondary elements that may be of importance to the development of evidence-based responses will also be included. Using robust search, retrieval, and methodological approaches to minimise potential sources of research bias with the proposed Evidence and Gap Map (EGM), we will summarise the existing and emerging evidence for the first time. The map will be made publicly available and interested parties will have access to a visual presentation of knowledge of the role of wildlife trade on human infectious diseases.

### Managed Reviews: NER To Determine Which Genes Matter

**Author:** Thomas Luechtefeld at Insilica LLC

**Sysrev:** https://sysrev.com/p/3144 - Gene Hunter NER

#### Description

Although there is an acknowledged nomenclature for genes, they may be described in the scientific literature in ways that have evolved over time. For example, *MLH1* may be written as “mutL homolog 1,” but previously may have been referred to as “COCA2”. These differences encumber rapid and accurate gene-phenotype associations.

Sysrev’s named entity recognition models and payment platform were combined to create a gene named entity recognition tool. In this project, users annotated genes in 10,000 sentences from medical abstracts from Pubmed (available at: sysrev.com/p/3144). In aggregate, 10 users were paid $1000, and the project was completed in 2 weeks. This data was used to create a named entity recognition model that could then automate extraction of genes from text.

### Education: Coursera Evidence-Based Toxicology

**Author:** Lena Smirnova at Center for Alternatives to Animal Testing at Johns Hopkins University, Bloomberg School of Public Health

**Sysrev:**

https://sysrev.com/p/3509 - Evidence-Based Toxicology Assignment 2018
https://sysrev.com/p/27474 - EBT Course Assignment Year 3
https://sysrev.com/p/29023 - GU systematic review 2019/2020

#### Description

Sysrev has value for educational purposes as a way to train individuals in SER practices. Dr. Lena Smirnova created an Evidence-Based Toxicology program on Coursera with a sysrev based assignment. Students were asked to apply SER principles in a final assessment that requires them to screen studies and resolve conflicts with other students according to pre-specified inclusion and exclusion criteria (available at: sysrev.com/p/3509). Over 411 students have now completed the course and thousands of labels have been assigned in the sysrevs. These courses indicate the potential for Sysrev to be used in teaching while simultaneously organizing valuable data from thousands of documents. The ease with which reviewers can navigate Sysrev and its alignment to FAIR principles make it an ideal choice for educational purposes.

### Limited Understanding Of Bushfire Impacts On Australian Invertebrates [25]

**Author:** Manu E. Saunders^1^, Philip S. Barton^1,2^, James R. M. Bickerstaff^3^, Lindsey Frost^1^, Tanya Latty^4^, Bryan D. Lessard^5^, Elizabeth C. Lowe^6^, Juanita Rodriguez^5^, Thomas E. White^4^, Kate D. L. Umbers^3,7^

1. School of Environmental and Rural Sciences, University of New England, Armidale, NSW, Australia.
2. School of Science, Psychology and Sport, Federation University Australia, Mount Helen, VC, Australia.
3. Hawkesbury Institute for the Environment, Western Sydney University, Penrith, NSW, Australia.
4. School of Life and Environmental Sciences, University of Sydney, Sydney, NSW, Australia.
5. Australian National Insect Collection, CSIRO, Canberra, ACT, Australia.
6. Department of Biological Sciences, Macquarie University, Sydney, NSW, Australia.
7. School of Science, Western Sydney University, Penrith, NSW, Australia

**Sysrev:** https://sysrev.com/p/24557- Fire and Australian invertebrates

#### Description

After Australia’s 2019-20 catastrophic bushfire disaster, media coverage and official estimates of biodiversity loss, as well as discussion about priorities for post-fire conservation activities, were focused on vertebrate animals. Invertebrates, which make up 95% of all animals on Earth, were largely neglected in these discussions, except for a few charismatic species.

Identifying the gaps and opportunities in current knowledge is an important first step to design effective policies that support biodiversity conservation and ecosystem recovery. Despite the ecological importance of invertebrates, and their taxonomic dominance of the animals, there are very few general literature reviews of fire impacts on invertebrates, and most are decades old or specific to a geographic location. This means that we have very little knowledge available that is relevant to contemporary global climate conditions, or Australian ecosystems generally.

To understand existing knowledge, we synthesised the published literature using a standardised review method to find out what evidence is available to inform invertebrate conservation policies in the age of extreme fire events. We used a structured search in Scopus to collate peer-reviewed studies that measured the impacts of fire on any group of invertebrates in an Australian ecosystem. We then used Sysrev to screen studies via abstracts, before collating the final set of studies for manual data extraction.

Our results show that there is very little awareness of how fire affects invertebrate communities in Australian ecosystems. Peer reviewed studies were available for only six of the more than 30 invertebrate phyla and 88% of targeted studies were on arthropods, predominantly ants. Nearly all studies (94%) were conducted on land, with only four studies measuring impacts in freshwater habitats. We found no studies of impacts on marine invertebrates in Australian waters, which is concerning – ash and post-fire runoff can have major effects on marine ecosystems, yet we didn’t find any evidence of how this is affecting marine invertebrates. This is a real problem because fire risk and severity is increasing globally. For effective climate change and disaster recovery policies, we need to know how to identify key risks and opportunities for ecosystem recovery and biodiversity conservation in the face of these new challenging conditions.

### Pre-clinical and Translational Studies on Macrophage Polarization in Nanoparticle-based Cancer Immunotherapy

**Author:** Colette Bilynsky, Wonhee Han, Anushree Gupta, Dasia Aldarondo, Hannah Fox, Chloe Brown, Archippe Mbembo, Megha Anand, Marissa McAfee, Jacob Bauldock, Melanie Gainey, Sarah Young and Elizabeth Wayne at Carnegie Mellon University

**Sysrev**

https://sysrev.com/p/31994 - Phase 1 title/abstract screening
https://sysrev.com/p/37476 - Phase 2 title/abstract
https://sysrev.com/p/43536 - Full text screening

#### Description

The protocol for this project is registered on the Open Science Framework [26].

The recent recognition of cancer as an inherently immunological disease has led to interesting opportunities for immunotherapy development. Tumors thrive by promoting an immunosuppressive environment, upregulating non-malignant stromal cells that support tumor proliferation and migration and downregulating those that might support a cytotoxic immune response. Tumor-associated macrophages (TAMs) are an important component of the tumor microenvironment, comprising up to 50% of the cells in solid tumors [27]. When TAMs are activated towards M2 polarization, which is anti-inflammatory and pro-tumoral, they contribute to tumor progression and indicate worse patient survival. Many therapy strategies have attempted to alter the polarization of macrophages’ “re-programming” or prevent monocyte tumor infiltration [28].

Nanoparticles are widely used to enhance the ability to target macrophages within tumor environments. Nanoparticles enable the customization of particle chemistries to modulate body pharmacokinetics, the encapsulation of multiple drugs or drugs that are water insoluble, and allows tunable control of drug release rates. However, nanoparticle customizability creates an exponential number of potential therapy design strategies. Thousands of studies have been published on the application of nanotechnology in cancer treatment and diagnosis but there have only been a very small number of FDA approved nanotherapeutics [29]. This translational gap suggests that there is much to understand between nano-formulations of cancer therapeutics that are falling short of clinical implementation.

The purpose of this scoping review is to utilize data taken from studies implementing nanomedicine for cancer treatment and diagnosis and determine how these formulations modulate tumor-associated macrophage polarization. Specifically, we seek to answer whether there is an identifiable relationship between nanoparticle characterization (i.e. type, size, shape charge, cargo) and their ability to modulate macrophage polarization in the tumor microenvironment. This will allow us to assess the field of cancer therapeutics to understand the effectiveness of nano-therapeutics in an immunological context. This work is unique in its field as a comprehensive, systematic scoping review of how cancer nanotherapeutics interact with macrophages. Some previously published systematic reviews in cancer nanomedicine have focused on specific subsections, like using nanoparticles for actively-targeted delivery of chemotherapeutics or herbal medicine [30]. Meanwhile, systematic reviews in cancer research have recently increased in frequency. However, these reviews mostly focus on clinical data with patients and not on research using in vitro or animal models of cancer [31]. Thus, our scoping review has a role in the field as a systematic overview of cancer nanotherapeutics to provide understanding of how these formulations affect drug development and efficacy in treating cancer.

Sysrev has proven to be an excellent platform for managing the screening of thousands of studies by a large collaborative team new to the evidence synthesis process. Moreover, we have taken advantage of Sysrev’s machine learning capabilities to automatically exclude a large number of articles using the predictions made based on the manual inclusion and exclusion of studies. This has greatly reduced the time and effort required to get through the title/abstract screening phase of the project allowing us to proceed more quickly to full text screening and data extraction.

### Indigenous Knowledge On Climate Change Adaptation: A Global Evidence Map Of Academic Literature

**Author:** Jan Petzold^1^, Nadine Andrews^2^, James D. Ford^3^, Christopher Hedemann^4^, Julio C. Postigo^4^

1. Center for Earth System Research and Sustainability, University of Hamburg, Hamburg, Germany.
2. The Pentland Centre for Sustainability in Business, Lancaster University, Lancaster, United Kingdom.
3. Priestley International Centre for Climate, University of Leeds, Leeds, United Kingdom.
4. Department of Geography, Indiana University Bloomington, Bloomington, United States of America

**Sysrev:** private project

#### Description

There is emerging evidence of the important role of indigenous knowledge for climate change adaptation. The necessity to consider different knowledge systems in climate change research has been established in the fifth assessment report (AR5) of the Intergovernmental Panel on Climate Change (IPCC). However, gaps in author expertise and inconsistent assessment by the IPCC lead to a regionally heterogeneous and thematically generic coverage of the topic. We conducted a scoping review of peer-reviewed academic literature to support better integration of the existing and emerging research on indigenous knowledge in IPCC assessments. The research question underpinning this scoping review is: How is evidence of indigenous knowledge on climate change adaptation geographically and thematically distributed in the peer-reviewed academic literature?

As the first systematic global evidence map of indigenous knowledge in the climate adaptation literature, the study provides an overview of the evidence of indigenous knowledge for adaptation across regions and categorises relevant concepts related to indigenous knowledge and their contexts in the climate change literature across disciplines. The results show knowledge clusters around tropical rural areas, subtropics, drylands, and adaptation through planning and practice and behavioural measures. Knowledge gaps include research in northern and central Africa, northern Asia, South America, Australia, urban areas, and adaptation through capacity building, as well as institutional and psychological adaptation. This review supports the assessment of indigenous knowledge in the IPCC AR6 and also provides a basis for follow-up research, e.g., bibliometric analysis, primary research of underrepresented regions, and review of grey literature.

### Synthesized evidence on conflict and war: a systematic map and critical appraisal of systematic reviews

**Author:** Zahra Saad^**1**^, Tamara Lotfi^**1**^, Hussein Ismail^**2**^, Neal Haddaway^**3**^, Elie Akl^**1**^

**1** The Global Evidence Synthesis Initiative (GESI) Secretariat, American University of Beirut (AUB), Lebanon. **2** American University of Beirut, Lebanon. **3** Stockholm Environment Institute, Stockholm

**Sysrev**

https://sysrev.com/p/39400 - Example of critical appraisal
https://sysrev.com/p/36852 - Example of data abstraction

#### Description

The Global Evidence Synthesis Initiative (GESI) Secretariat is developing an evidence gap map of published evidence synthesis in the humanitarian field. The objective of the systematic mapping is to:

1. rigorously identify and critically appraise published systematic reviews in the humanitarian field, specifically on conflict and war.
2. publish a user-friendly, interactive and readily digestible map of the evidence accessible to stakeholders including humanitarian practitioners, policy makers, and researchers.

The team developed search strategies for 11 databases, and then completed title and abstract screening for 3214 records. The full text screening of 601 eligible papers led to the inclusion of 340 evidence synthesis products. Then, the GESI Secretariat invited the GESI Network members to take part in data abstraction and critical appraisal as a collaborative approach using the free online software Sysrev.

The GESI Secretariat connected with the Sysrev team and presented an introductory webinar to train the participants on how to use the platform. To split the work, the participants were divided into groups of 2 reviewers each. On Sysrev, 12 cloned data abstraction projects of twenty-four variables were created and 8 cloned critical appraisal projects of 11 variables were formed, example links for access to these public projects are given above. The software offered a feature to view discrepancies in the answers allowing reviewers working in pairs to discuss them and accordingly resolve disagreements. At a later stage, the admin of the project exported all data in an excel format and then combined all results for data analysis. The team is currently developing the manuscript.

### EntoGEM: A Community-Driven Systematic Mapping Project To Build A Global Evidence Map Of Insect Population And Biodiversity Trends

**Author:** Eliza .M. Grames^1^, Graham.A. Montgomery^2^

1. University of Connecticut, Storrs USA 06351
2. University of California Los Angeles, Los Angeles USA 90095

**Sysrev:** https://sysrev.com/p/16612 - EntoGEM: a systematic map of global insect population and biodiversity trends

#### Description

EntoGEM is a community-driven project that aims to compile evidence about the current status and trends of global insect populations and biodiversity. Our goal is to build a systematic map database that contains all studies and datasets, published and unpublished, that contain evidence regarding long-term insect population and biodiversity declines, increases, or lack of changes over time. The final product will be publicly accessible, interactive, and searchable by metadata coded for each study, such as location, taxa, duration and time period of study, and other variables listed in the pre-registered protocol [32]. By identifying what studies and datasets exist, this project will help prioritize future research on insect conservation, guide granting by funding agencies, and facilitate follow-up syntheses.

Because the comprehensive search for the project yielded over 144,000 results, we are using SysRev to complete targeted subsets of the search results, which then feed back into topic models that help to prioritize future screening efforts by the community. Although only roughly 5% of the EntoGEM search results have been screened by the community so far, already we have identified more than one hundred datasets that contain 10 or more years of insect population and biodiversity data that have not been referenced in any reviews, meta-analyses, or syntheses on insect decline to date [33].

#### Funding

EntoGEM was the first winner of the Sysrev mini-grants, small grants to incentivize public health research. The project is also funded by the Doris Duke Charitable Foundation.

### SEAZIT - Systematic Evaluation Of The Application Of Zebrafish In Toxicology

**Author:** Kristin Ryan^**1**^, Jon Hamm^**2**^, Neepa Choksi^**2**^, Lauren Browning^**2**^ and Nicole Kleinstreuer^**3**^

1. National Institutes of Health/National Institute of Environmental Health Sciences/Division of the National Toxicology Program/Systems Toxicology
2. Integrated Laboratory Systems, LLC.
3. National Institutes of Health/National Institute of Environmental Health Sciences/Division of the National Toxicology Program/National Toxicology Program Interagency Center for the Evaluation of Alternative Toxicological Methods

**Sysrev**

https://sysrev.com/p/54429 - Amoxicillin Part 1
https://sysrev.com/p/55474 - Aspirin Part 1

22 other related reviews

#### Description

NICEATM is currently working on a project entitled Systematic Evaluation of the Application of Zebrafish in Toxicology (SEAZIT), aimed at understanding the utility of zebrafish for toxicological screening and the impact of key protocol variations on the data generated by this small model organism system. The current phase of SEAZIT involves an interlaboratory study utilizing 39 chemicals to compare several exposure conditions. Developmental toxicity results and outcomes available in published literature on these chemicals will be used to interpret the outcome of the interlaboratory study. Broad literature searches focused on retrieval of developmental toxicity studies in mammalian and zebrafish models have been conducted in PubMed. Relevant studies (5-10 per chemical) will be reviewed and used to assist in results interpretation from the proposed interlaboratory study.

#### Funding

This project was funded with federal funds from the NIEHS, NIH under Contract No. HHSN273201500010C

### Cardiotoxicity Evaluation

**Author:** Nicole Kleinstreuer^1^, Neepa Choksi^2^, Lauren Browning^2^, Amber Daniel^2^

1. National Institutes of Health/National Institute of Environmental Health Sciences/Division of the National Toxicology Program/National Toxicology Program Interagency Center for the Evaluation of Alternative Toxicological Methods
2. Integrated Laboratory Systems, LLC.

**Sysrev**

https://sysrev.com/p/32069 - In Vitro Cardiovascular Methods
https://sysrev.com/p/33691 - In Vitro Cardiovascular Methods – Predictions Subset

#### Description

Cardiotoxicity is a major cause of failure of new drugs in mid- to late-stage development, cardiovascular disease is one of the leading causes of mortality in the US and worldwide. While cardiovascular health effects of several environmental exposures such as smoking, air pollution, and lead are well characterized, exposure to other environmental chemicals represents another potentially significant but underappreciated risk factor contributing to the development and severity of cardiovascular disease, and reliable methods to detect potential chemical cardiotoxicity need to be identified and applied. Most cardiotoxicity testing is currently done in animals, but these tests are expensive, time-consuming, and can fail to fully predict effects in humans. NICEATM and NTP staff used Sysrev.com to conduct literature searches to identify in vitro methods with mechanistic targets that can be mapped to one of six identified cardiotoxicity “failure modes” [34]. These methods could be used to develop integrated testing strategies that are predictive of cardiovascular toxicity and provide human-relevant mechanistic information.

#### Funding

This project was funded with federal funds from the NIEHS, NIH under Contract No. HHSN273201500010C.

We are grateful to the many teams that submitted case studies. There have now been thousands of public projects on Sysrev, and we were unable to request case studies from all project owners. Other projects can be discovered by using the Sysrev.com search functionality. Sysrev makes efforts to index all public projects and make them discoverable by major search engines and via the on site search function. For example, to find projects, users, and groups related to “cancer” use the search https://sysrev.com/search?q=cancer&p=1&type=projects. This results in 35 projects.

## Meta-Analysis

Sysrev generates millions of data points for article labels, model predictions, and projects. Sysrev actively uses that data to improve the visibility of systematic review, build a review community, and improve the state of machine learning for human evidence review and data extraction.

### A Review Community

As of March 2021, 3224 sysrevs have been created on Sysrev.com, 442,265 documents have been reviewed, and 1.8 million review tasks have been completed. Sysrev aims to provide a platform where reviewer communities can easily form to develop and share their methods and data.

The force layout diagram in Figure 6A looks at 3 of the larger communities on sysrev.com. Each pink node is a user, and each green node is a sysrev. Connections are drawn between sysrevs and their members. Each of these authors and a selection of their projects are referenced in the case studies section.

**Figure 6.**
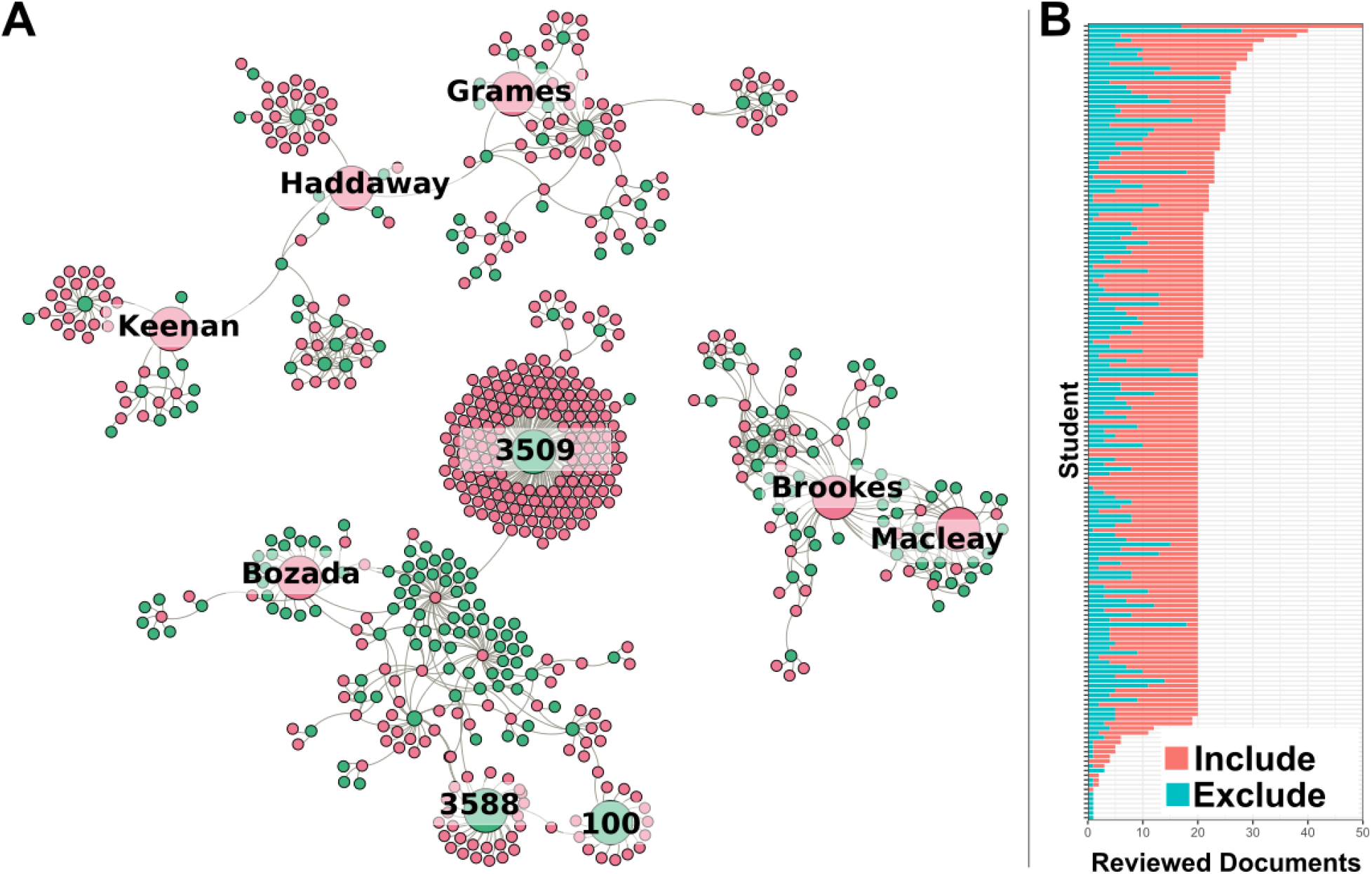
**A.** Three of the larger communities on sysrev.com. Pink nodes represent users, green nodes represent projects. Edges link projects with their users. Several notable projects and users have been enlarged for discussion. **B.** Bar chart representing completed document reviews for each student in sysrev.com/p/3509, an educational review focused on evidence based toxicology by Dr. Lena Smirnova. Each bar represents a student, students were asked to complete a review of 20 documents, blue bars indicate excluded articles, red bars indicate included articles in an article screening exercise.

Grames and Keenan were recipients of Sysrev mini-grants, which helped in small part to fund their research on insect population biodiversity trends (see case study) and chronic kidney disease in the context of Covid-19 (see case study). Both authors collaborated with Dr.

Haddaway (referenced in several case studies), and altogether the three have spawned multiple reviewer communities.

Bozada is a sysrev.com employee who has created many projects (public and private) for managed reviews. There is a chain of reviewer / sysrev membership links that lead to sysrev.com/p/3588 and sysrev.com/p/100 which are projects completed by toxicology and cancer research groups, the latter of which represents a hepatotoxicity project by the Evidence-Based Toxicology Collaboration led by Dr. Tsaioun at Johns Hopkins which provided support for a report on “Performance of preclinical models in predicting drug-induced liver injury in humans: a systematic review” [35].

Figure 1B Shows the progression of public project sysrev.com/p/3509. This is an educational project teaching students to review toxicology data. It is an assignment of a coursera course on evidence based toxicology by Dr. Smirnova at Johns Hopkins School of Public Health. As of March 1st 2021, over 170 students have extracted 24,791 label answers from 1,581 zebrafish toxicology documents, making this one of Sysrev’s largest public projects by number of users.

### Active Learning To Accelerate Review

Sysrev uses active learning to train screening algorithms and prioritize reviews. The method Sysrev uses is to first distribute random articles for review, then train a model on reviewer answers, then distribute articles with the highest prediction entropy for inclusion (closest to 0.5 probability for inclusion), then retrain models, and repeat the process. The intention of active learning is to accelerate machine learning, so as to build accurate models as quickly as possible and minimize the use of reviewer times on easily modeled documents.

Figure 7A evaluates the normalized log score of models as a function of the number of articles that are reviewed before, and are subsequently used in model training. The normalized log score is calculated on the last 20% of reviewed articles (articles that have not been used in the training set for any of the visualized models). 64 projects were chosen for this analysis that had at least 1000 reviewed articles, and had a minimum of 20% included articles and 20% excluded articles. This was done because Sysrev model accuracy and balanced accuracy suffers in projects with a highly imbalanced inclusion/exclusion ratio. Articles that were discordant (receiving an include from one reviewer and an exclude from a different reviewer) were discarded. The formula used for normalized score was:

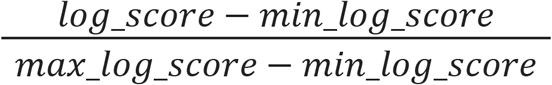

**Figure 7.**
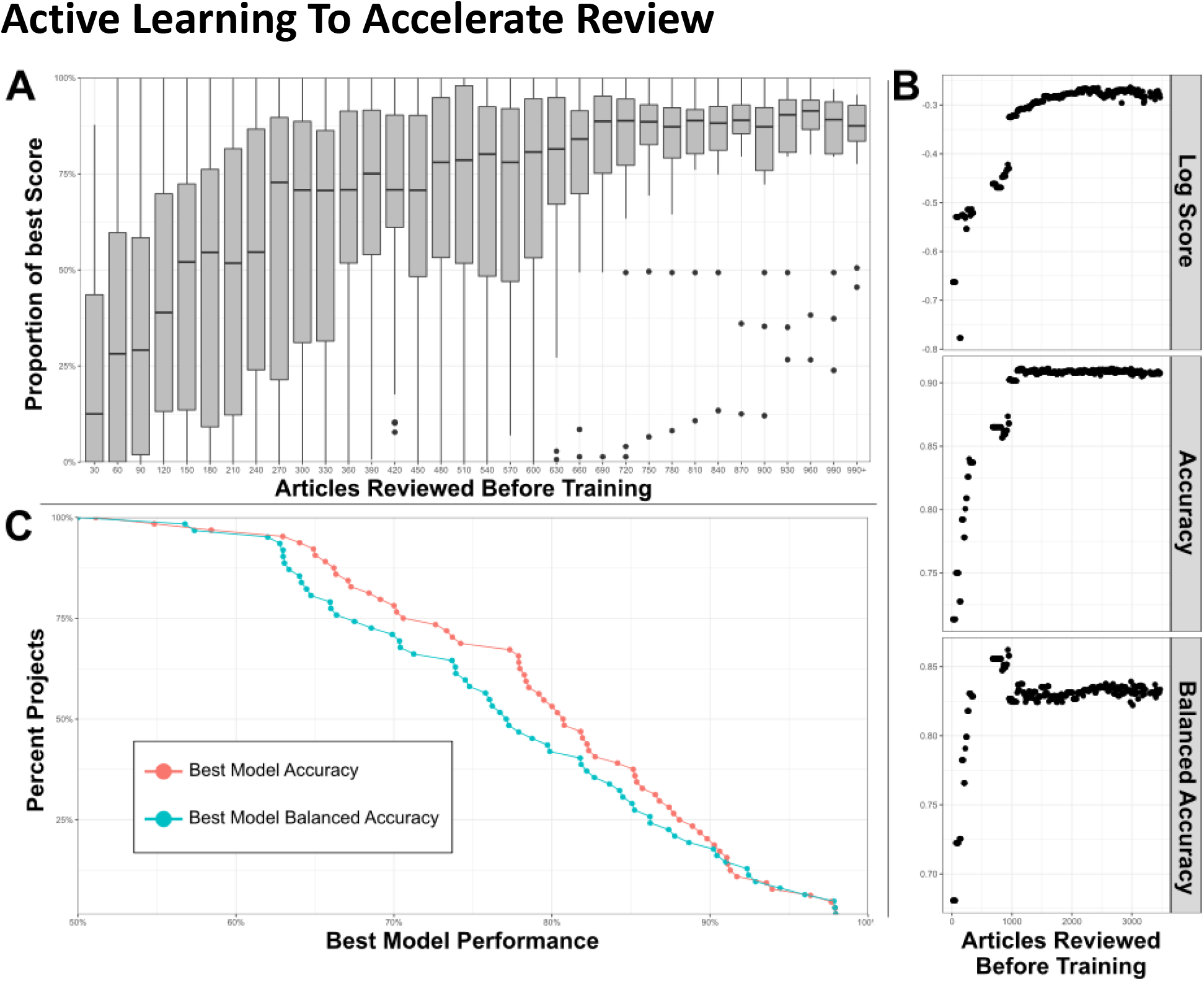
**A.** Box plots indicate distribution of model performance relative to the worst and best model in a given model’s project. Models are bucketed according to the number of articles labeled before model training. Models improve rapidly until 300 articles have been reviewed. **B.** Accuracy metrics for a large sysrev reviewing insect population changes. Model performance is charted as a function of number of articles used in training, across 3 performance metrics, and evaluated on a consistent holdout set. **C.** Best model accuracy and balanced accuracy evaluated in 64 sysrevs.

Log score is the logarithmic scoring rule for evaluating prediction models, this is a proper scoring rule that assigns greater values to models that assign high confidence to correct predictions, and low confidence to incorrect predictions [36]. In this case, log score is maximized (with a value of 0) by models that assign a 1.0 prediction to articles that were concordantly reviewed as inclusion articles and a 0.0 prediction to articles that were concordantly excluded. An additional normalization method of subtracting the log score of the worst performing model in a project and dividing by the difference between the log score of the best and worst model in a project transforms this metric into a proportion receiving a value of 0.0 for the worst models in a project and a value of 1.0 for the best. Values closer to 1.0 indicate model performance close to the best achieved in a project. This normalization method is used to make models with different baseline performance more comparable and allow for an aggregate evaluation of the number of articles required to build models close to maximum performance in a sysrev.

Figure 7A provides a boxplots visualizing inclusion model performance relative to the best and worst inclusion model for a project, a value of 100% indicates that the model has log score performance equal to the best model for a project, 0% indicates a model with log score equal to the worst model for the project. The median performance (relative to the best project model) of the first models built in projects (after a review of just 30 articles) is close to 0% for most projects. After reviewing 690 articles, most models are relatively close to the best model that will ever be achieved for their projects. There is no guarantee that models will perform well after reviewing 690 articles, actual performance is dependent on many factors. This figure simply demonstrates that the Sysrev model generating algorithm usually converges to some maximum performance for the average sysrev after a few hundred articles have been reviewed. In some cases this convergence occurs much earlier, and in some cases it occurs much later. One of the goals of active learning is to accelerate the pace at which maximum model performance is achieved.

Inclusion model performance metrics are visualized as a function of articles reviewed before model training for the entogem sysrev (see case study and sysrev.com/p/16612) in Figure 7B. Dr. Grames and her team reviewed 7091 articles, the last 20% of articles were used to evaluate models, ~1500 discordant articles were not included in this analysis. Model accuracy reaches a maximum after ~1000 concordant articles have been reviewed, model balanced accuracy actually declines slightly to 82.5% at the 1000 article point. Log score continues to improve for several thousand more article reviews. This indicates that, for this project, the models continue to improve by generating higher confidence correct predictions and lower confidence incorrect predictions, with only small changes in the total number of incorrect and correct predictions. Models with high confidence of correct predictions, and low confidence of incorrect predictions are more valuable than otherwise equally accurate models.

While proper scoring rules like log score provide a good means for comparing model performance, sometimes it is easier to interpret accuracy (number of correct predictions divided by total predictions) and balanced accuracy (an average of sensitivity and specificity, correctly predicted inclusions and exclusions in this case). Figure 7C provides a cumulative distribution of the best model accuracy and best model balanced accuracy aggregated over projects. Of the 64 surveyed projects, 75% had a balanced accuracy and accuracy of 68% or higher, 50% of these projects accurately predicted concordant human screening decisions 80% of the time or higher. None of the models achieved perfect performance. It is our belief (untested in this paper), that Sysrev screening model performance is strongly dependent on the types of articles being reviewed, on the reviewers performing the project, on the complexity of inclusion criteria, on statistical factors like the balance of included articles to excluded articles and many other factors. There is no guarantee that Sysrev models will perform well on a given project and Sysrev provides tools to evaluate model performance during project progression to allow administrators to determine the value of these tools on a project by project basis. Nevertheless, it is informative to observe that, in aggregate, these models have performed well for many sysrevs. This is a difficult task that indicates that the sysrev model architecture and training method generalizes well across many different kinds of projects. Future research will provide more information on sysrev modeling efforts.

## Conclusion

We developed Sysrev to be a FAIR-compliant, discipline-agnostic platform for data curation and SERs. Sysrev can be easily adopted for non-biomedical research, such as environmental, climate, or industrial purposes. Existing datasets are often proprietary or have limited applicability beyond their original purpose, which contributes to the promulgation of data silos. Importantly, much of the data captured in both primary research and SERs is published in a manner that inhibits the application of the data to be re-used for new investigations. This is especially problematic for investigations where users may not be aware of relevant data from cross-discipline/industry studies. Sysrev’s public projects are accessible, clonable, and transparent to encourage the responsible reuse of data and support broad research goals. Sysrev has been used for thousands of review and data curation projects. Users agree that Sysrev makes the SER process run more efficiently, even with large teams of more than 100 users or those with tens of thousands of studies/data. Further improvements to improve the user experience are planned and in development. Sysrev offers several access options, including a free version with unlimited publicly available projects or pro/enterprise versions that include additional functionality. Sysrev is privately funded by insilica.co and can be accessed at https://sysrev.com.

## References

1. Burns PB, Rohrich RJ, Chung KC (2011) The Levels of Evidence and Their Role in Evidence-Based Medicine. Plast Reconstr Surg 128:305–310. https://doi.org/10.1097/prs.0b013e318219c171

2. Mallett R, Hagen-Zanker J, Slater R, Duvendack M (2012) The benefits and challenges of using systematic reviews in international development research. J Dev Effect 4:445–455. https://doi.org/10.1080/19439342.2012.711342

3. Mierden SV der, Tsaioun K, Bleich A, Leenaars CHC (2019) Software tools for literature screening in systematic reviews in biomedical research. Altex. https://doi.org/10.14573/altex.1902131

4. Gates A, Guitard S, Pillay J, Elliott SA, Dyson MP, Newton AS, Hartling L (2019) Performance and usability of machine learning for screening in systematic reviews: a comparative evaluation of three tools. Syst Rev 8:278. https://doi.org/10.1186/s13643-019-1222-2

5. Harrison H, Griffin SJ, Kuhn I, Usher-Smith JA (2020) Software tools to support title and abstract screening for systematic reviews in healthcare: an evaluation. Bmc Med Res Methodol 20:7. https://doi.org/10.1186/s12874-020-0897-3

6. Hariri RH, Fredericks EM, Bowers KM (2019) Uncertainty in big data analytics: survey, opportunities, and challenges. J Big Data 6:44. https://doi.org/10.1186/s40537-019-0206-3

7. Wilkinson MD, Dumontier M, Aalbersberg IjJ, Appleton G, Axton M, Baak A, Blomberg N, Boiten J-W, Santos LB da S, Bourne PE, Bouwman J, Brookes AJ, Clark T, Crosas M, Dillo I, Dumon O, Edmunds S, Evelo CT, Finkers R, Gonzalez-Beltran A, Gray AJG, Groth P, Goble C, Grethe JS, Heringa J, Hoen PAC’t, Hooft R, Kuhn T, Kok R, Kok J, Lusher SJ, Martone ME, Mons A, Packer AL, Persson B, Rocca-Serra P, Roos M, Schaik R van, Sansone S-A, Schultes E, Sengstag T, Slater T, Strawn G, Swertz MA, Thompson M, Lei J van der, Mulligen E van, Velterop J, Waagmeester A, Wittenburg P, Wolstencroft K, Zhao J, Mons B (2016) The FAIR Guiding Principles for scientific data management and stewardship. Sci Data 3:160018. https://doi.org/10.1038/sdata.2016.18

8. Wallis JC, Rolando E, Borgman CL (2013) If We Share Data, Will Anyone Use Them? Data Sharing and Reuse in the Long Tail of Science and Technology. Plos One 8:e67332. https://doi.org/10.1371/journal.pone.0067332

9. Heidorn PB (2008) Shedding Light on the Dark Data in the Long Tail of Science. Libr Trends 57:280–299. https://doi.org/10.1353/lib.0.0036

10. Tryka KA, Hao L, Sturcke A, Jin Y, Wang ZY, Ziyabari L, Lee M, Popova N, Sharopova N, Kimura M, Feolo M (2014) NCBI’s Database of Genotypes and Phenotypes: dbGaP. Nucleic Acids Res 42:D975–D979. https://doi.org/10.1093/nar/gkt1211

11. Arnaud E, Laporte M-A, Kim S, Aubert C, Leonelli S, Miro B, Cooper L, Jaiswal P, Kruseman G, Shrestha R, Buttigieg PL, Mungall CJ, Pietragalla J, Agbona A, Muliro J, Detras J, Hualla V, Rathore A, Das RR, Dieng I, Bauchet G, Menda N, Pommier C, Shaw F, Lyon D, Mwanzia L, Juarez H, Bonaiuti E, Chiputwa B, Obileye O, Auzoux S, Yeumo ED, Mueller LA, Silverstein K, Lafargue A, Antezana E, Devare M, King B (2020) The Ontologies Community of Practice: A CGIAR Initiative for Big Data in Agrifood Systems. Patterns 1:100105. https://doi.org/10.1016/j.patter.2020.100105

12. David E, Madec S, Sadeghi-Tehran P, Aasen H, Zheng B, Liu S, Kirchgessner N, Ishikawa G, Nagasawa K, Badhon MA, Pozniak C, Solan B de, Hund A, Chapman SC, Baret F, Stavness I, Guo W (2020) Global Wheat Head Detection (GWHD) Dataset: A Large and Diverse Dataset of High-Resolution RGB-Labelled Images to Develop and Benchmark Wheat Head Detection Methods. Plant Phenomics 2020:1–12. https://doi.org/10.34133/2020/3521852

13. Bozada T (2020) What is Sysrev for? Literature Reviews and Data Curation. https://blog.sysrev.com/literature-review-data-curation/

14. Settles B (2009) Active learning literature survey

15. Yang B, Sun J-T, Wang T, Chen Z (2009) Effective multi-label active learning for text classification. 917–926. https://doi.org/10.1145/1557019.1557119

16. Paller C, Tamashiro T, Luechtefeld T, Gravell A, Levine M (2020) Vitamin C and Cancer An Overview of Recent Clinical Trials. In: Vissers QC and MCM (ed) Cancer and Vitamin C, 1st Edition. p 64

17. Kilgore WW (1990) California’s proposition 65: Extrapolating animal toxicity to humans. Am J Ind Med 18:491–494. https://doi.org/10.1002/ajim.4700180422

18. Luechtefeld T (2019) Sysrev Helps Create Chemical Transparency for Manufacturers. https://blog.sysrev.com/srg-sysrev-chemical-transparency/. Accessed 16 Mar 2021

19. Bozada TJ (2020) Supporting COVID Research: Rapid Reviews on Sysrev,. https://blog.sysrev.com/covid-rapid-review/. Accessed 16 Mar 2021

20. Mahalingasivam V, Craik A, Tomlinson LA, Ge L, Hou L, Wang Q, Yang K, Fogarty DG, Keenan C (2021) A Systematic Review of COVID-19 and Kidney Transplantation. Kidney Int Reports 6:24–45. https://doi.org/10.1016/j.ekir.2020.10.023

21. Sterne JA, Hernán MA, Reeves BC, Savović J, Berkman ND, Viswanathan M, Henry D, Altman DG, Ansari MT, Boutron I, Carpenter JR, Chan A-W, Churchill R, Deeks JJ, Hróbjartsson A, Kirkham J, Jüni P, Loke YK, Pigott TD, Ramsay CR, Regidor D, Rothstein HR, Sandhu L, Santaguida PL, Schünemann HJ, Shea B, Shrier I, Tugwell P, Turner L, Valentine JC, Waddington H, Waters E, Wells GA, Whiting PF, Higgins JP (2016) ROBINS-I: a tool for assessing risk of bias in non-randomised studies of interventions. Bmj 355:i4919. https://doi.org/10.1136/bmj.i4919

22. UNEP, ILRI (2020) Preventing the Next Pandemic: Zoonotic diseases and how to break the chain of transmission.

23. Daszak P, Neves C das, Amuasi J, Haymen D, Kuiken T, Roche B, Zambrana-Torrelio C, Buss P, Dundarova H, Feferholtz Y, others (2020) Workshop Report on Biodiversity and Pandemics of the Intergovernmental Platform on Biodiversity and Ecosystem Services

24. Cáceres-Escobar H, Broad S, Challender DWS, Kinnaird MF, Keenan C, Lichtenstein G, Rodriguez JP, Roe D, Rouan AD, Dijk PPV, al. et (2020) PROTOCOL: An Evidence and Gap Map (EGM) about the Roles and Risks of Wildlife Trade in the Emergence of Human Infectious Diseases and Zoonoses. https://doi.org/10.6084/m9.figshare.13392275.v2

25. Saunders ME, Barton PS, Bickerstaff JRM, Frost L, Latty T, Lessard BD, Lowe EC, Rodriguez J, White TE, Umbers KDL (2021) Limited understanding of bushfire impacts on Australian invertebrates. Insect Conservation and Diversity. https://doi.org/https://doi.org/10.1111/icad.12493

26. Bilynsky C, Han W, Gupta A, Aldarondo D, Fox H, Brown C, Mbembo A, Anand M, McAfee M, Bauldock J, Gainey M, Young S, Wayne E (2021) Scoping Review of Pre-clinical and Translational Studies on Macrophage Polarization in Nanoparticle-based Cancer Immunotherapy. https://doi.org/https://doi.org/10.17605/OSF.IO/HWD2R

27. Zhou J, Tang Z, Gao S, Li C, Feng Y, Zhou X (2020) Tumor-Associated Macrophages: Recent Insights and Therapies. Frontiers Oncol 10:188. https://doi.org/10.3389/fonc.2020.00188

28. Noy R, Pollard JW (2014) Tumor-Associated Macrophages: From Mechanisms to Therapy. Immunity 41:49–61. https://doi.org/10.1016/j.immuni.2014.06.010

29. Salvioni L, Rizzuto MA, Bertolini JA, Pandolfi L, Colombo M, Prosperi D (2019) Thirty Years of Cancer Nanomedicine: Success, Frustration, and Hope. Cancers 11:1855. https://doi.org/10.3390/cancers11121855

30. Muhamad N, Plengsuriyakarn T, Na-Bangchang K (2018) Application of active targeting nanoparticle delivery system for chemotherapeutic drugs and traditional/herbal medicines in cancer therapy: a systematic review. Int J Nanomed 13:3921–3935. https://doi.org/10.2147/ijn.s165210

31. Kelley GA, Kelley KS (2018) Systematic reviews and cancer research: a suggested stepwise approach. Bmc Cancer 18:246. https://doi.org/10.1186/s12885-018-4163-6

32. Grames E, Montgomery GA, Haddaway NR, Dicks LV, Elphick CS, Matson TA, Nakagawa S, Saunders ME, Tingley MW, White TE (2019) Trends in global insect abundance and biodiversity: A community-driven systematic map protocol. https://doi.org/10.17605/osf.io/q63uy

33. Grames E (2020) Preliminary results from the EntoGEM project May 06, 2020. https://doi.org/10.5281/zenodo.3814732

34. Krishna S, Berridge B, Kleinstreuer N (2020) High-Throughput Screening to Identify Chemical Cardiotoxic Potential. Chem Res Toxicol 34:566–583. https://doi.org/10.1021/acs.chemrestox.0c00382

35. Dirven H, Vist GE, Bandhakavi S, Mehta J, Fitch SE, Pound P, Ram R, Kincaid B, Leenaars CHC, Chen M, Wright RA, Tsaioun K (2021) Performance of preclinical models in predicting drug-induced liver injury in humans: a systematic review. Sci Rep-uk 11:6403. https://doi.org/10.1038/s41598-021-85708-2

36. Bickel JE (2007) Some Comparisons among Quadratic, Spherical, and Logarithmic Scoring Rules. Decis Anal 4:49–65. https://doi.org/10.1287/deca.1070.0089

